## Supplementary figures and images for "ATG5 is instrumental in the transition from autophagy to apoptosis during the degeneration of tick salivary glands"

### Supplemental Figure1

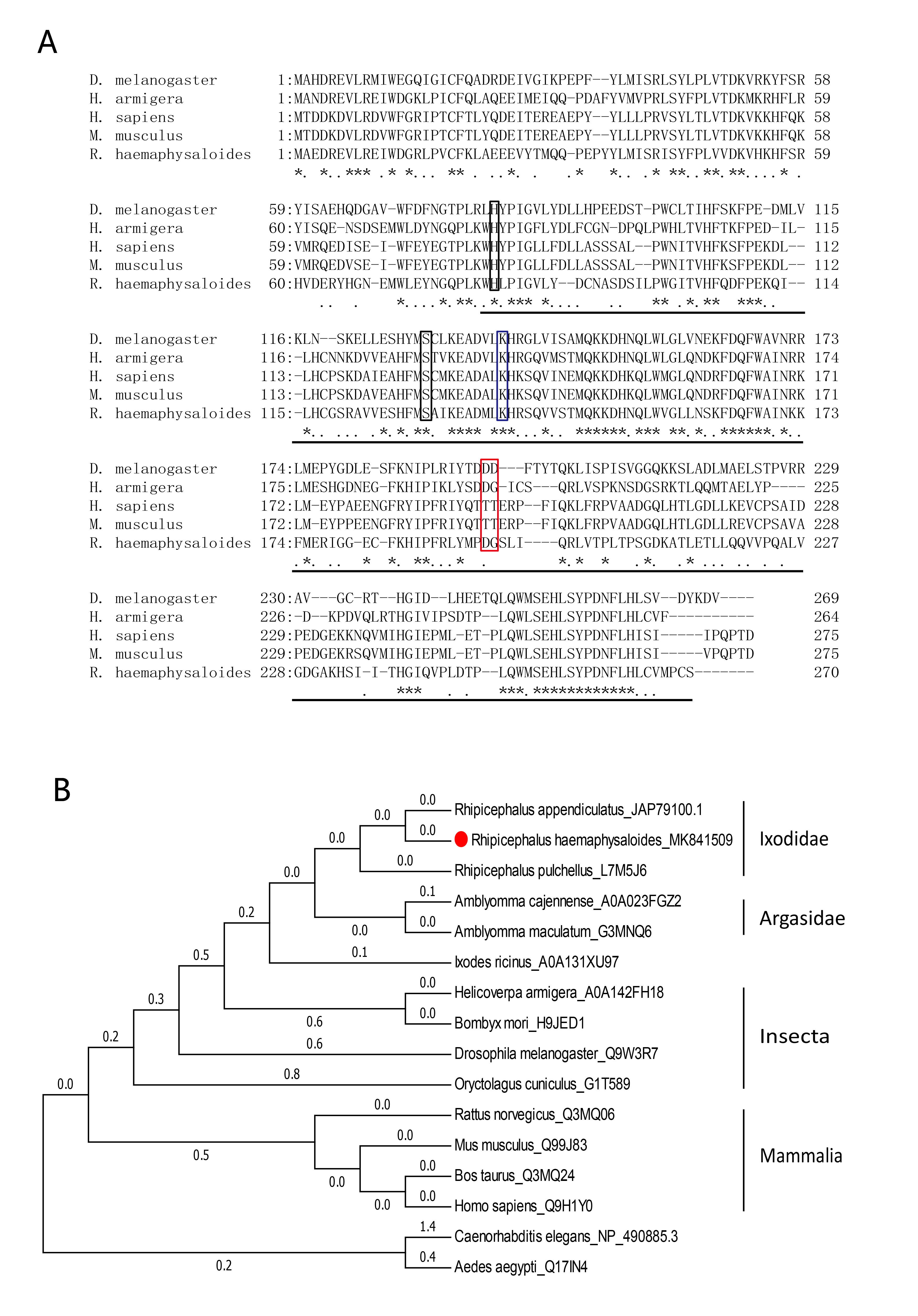
