## Supplemental Table 1 for "ATG5 is instrumental in the transition from autophagy to apoptosis during the degeneration of tick salivary glands"

**Supplement Table 1** Primers used for quantitative real-time polymerase chain reactions (qPCRs).

| **Primer name** | **Primer sequence** |
| --- | --- |
| ELF1A-S | CGTCTACAAGATTGGTGGCATT |
| ELF1A-A | CTCAGTGGTCAGGT TGGCAG |
| RhATG3-S | AGGTGACGAGGATGGCGGCTGGGTT |
| RhATG3-A | ATCCGCATCCAGCAGTCCACTCTCC |
| RhATG4B-S | CCTACCGCAAGAACTTCCCAGCAA |
| RhATG4B-A | TGCCAGTCTTTCCCAAGGTGCCGTC |
| RhATG4D-S | GGCAAGCAGGCGGGTGACTGGTATG |
| RhATG4D-A | TGCCCTTGACGCACGGAATGTATGT |
| RhATG5-S | GAAGCATCGCAGCCAAGTGGTCAGC |
| RhATG5-A | TGTGTGATGATGCTGTGTTTGGCTC |
| RhATG6-S | ACCTGATGTGCCTGGAAGACCCGAC |
| RhATG6-A | GGTAGGGAAGGCAGAAGTTGGAGTC |
| RhATG7-S | TGCCTGAGGAAGTGACCCTTGCCA |
| RhATG7-A | CCCGTTGGCACGACAAGAAAGGAG |
| RhATG8-S | CGCACGCAGTCAACGATAAGCAAGC |
| RhATG8-A | GGTATCTTTGATGGGAACCGTTGCCTG |
| RhATG9-S | CAGTGGGTGCGGAGGTGTCG |
| RhATG9-A | CCGTCCTCACCACTGCCTTCCT |
| RhATG10-S | AGAGCACTGTCTCCGTCATTTGTTC |
| RhATG10-A | TGGAAGTAGGGGACGCCTTGGAT |
| RhATG12-S | CCCCCGAGAAAACCGAAGCCCG |
| RhATG12-A | TTGGCTTCAACCCCAGGCATGCG |
| RhATG13-S | GGGAACCCTGACCCTCGGAGTGGA |
| RhATG13-A | GCATCCGCATACCCCGAGGGCTGA |
| RhATG14-F | GGGAAGCCATACAACGCACAAGGGA |
| RhATG14-R | GATGTCACGAGCGAGGTCTCCCACG |
| RhATG16-S | GCGGCTGCGGCACACACT |
| RhATG16-A | AATCTCGTTGGCACTGCTCTCTG |
| RhCaspase1-S | GATGCTGACAGTGGTGTGCCGCC |
| RhCaspase1-A | CCTGGTTTCGCCCTGAAGTAGAC |
| RhCaspase-7-S | CTCAGCGAGCGAAGGGGCACGGACA |
| RhCaspase-7-A | AGCAGACTGGGACAGACATCTCCGT |
| RhCaspase8-S | CGCCACAGTTTGGGACCACAGGA |
| RhCaspase8-A | CTTCGCCTTTCACCTGTGCCCCT |
| RhCaspase9-S | GCTGACAAGCCCACTGGCGAACAAC |
| RhCaspase9-A | CATTCAGAGCAGAGTCAGCAGTCCG |
| RhCalpain-1-S | CATCGTCCTGTCGCCAGTGGAGAAA |
| RhCalpain-1-A | CCGCCTTCGGACACGCCCAT |
| RhCalpain-2-S | CCTGGTCGGAGGCAGCGGCA |
| RhCalpain-2-A | CACCTGTGTGGCACCAGTGACGC |

**^a^**S, forward primer; A, reverse primer
