## Supplemental Table 2 for "ATG5 is instrumental in the transition from autophagy to apoptosis during the degeneration of tick salivary glands"

**Supplement Table 2** Primers for apoptosis and autophagy related gene cloning and vector construction

| **Primer name** | **Primer sequence** |
| --- | --- |
| RhATG5-S | CGCAGGTTCGATGTAAACATTTCTGTTC |
| RhATG5-A | AGACTAGCCGTAGAGTTTGAGAAATATCAC |
| RhATG5-pET-30a-S | GGCTGATATCGGATCCATGGCGGAAGATCGTGAAGTCTTAC |
| RhATG5-pET-30a-A | GTGCGGCCGCAAGCTTGGTTGTCTGGGTAACTGAGGTGTTCA |
| RhATG5^191-199Δ^-pET-30a-Sm | TTCAAGCACATACCTTTTCGCCTGAGGCTAGTCACGC |
| RhATG5^191-199Δ^-pET-30a-Am | GTGTCAATGGCGTGACTAGCCTCAGGCGAAAAGGTA |
| RhATG5 dsRNA-S1 | **GGATCCTAATACGACTCACTATAGG**CAAGGTCCATAAGCACTTCTCCAGG |
| RhATG5 dsRNA-A1 | CTCATCCATTGAAGCGGTGTGTCCA |
| RhATG5 dsRNA-S2 | CAAGGTCCATAAGCACTTCTCCAGG |
| RhATG5 dsRNA-A2 | **GGATCCTAATACGACTCACTATAGG**CTCATCCATTGAAGCGGTGTGTCCA |

**^a^**S, forward primer; A, reverse primer; Sm, forward primer for deletion mutation; Am, reverse primer for deletion mutation; the sequence in bold underlined indicates the sequence of the T7 promoter.
