## Supplemental Table 3 for "ATG5 is instrumental in the transition from autophagy to apoptosis during the degeneration of tick salivary glands"

**Supplement Table 3** Predicted epitope peptide sequences

| **Peptide name** | **Peptide sequence** |
| --- | --- |
| Rhcaspase7-1 | MAGMSGDDLQ |
| Rhcaspase7-2 | MEPFHGDVAV |
| RhATG8 | MKFQYKEEHPFEK |
